# CRISPR-Associated Transposases Enable Programmable DNA Integration in Plants

**DOI:** 10.64898/2026.07.15.738807

**Authors:** Yunqing Wang, Kimberley T. Muchenje, Ashot Papikian, Mark G. Legendre, Esther Leem, Gozde S. Demirer

## Abstract

Programmable DNA integration is a major challenge in plant genome engineering. CRISPR-associated transposases (CAST) catalyze efficient RNA-guided DNA integration without double-strand breaks, yet their activity has not been established in plants. Here, we reconstituted and engineered a Type I-F CAST for programmable DNA integration in plant cells. We validated expression of the wild-type Pseudoalteromonas CAST (PseCAST) machinery in plants and established targeted episomal integration in *Arabidopsis thaliana* protoplasts and chromosomal integration at a transgenic locus in *Nicotiana benthamiana*. The evolved PseCAST system, evoCAST, showed chromosomal integration efficiencies of 2.7%, representing a 6-fold improvement over wild-type PseCAST. evoCAST also enabled the insertion of cis-regulatory elements into a synthetic landing pad with 8% efficiency. evoCAST was subsequently retargeted to six endogenous genomic loci, demonstrating programmable integration across diverse chromosomal contexts. Finally, a cofactor screen identified the chromatin-associated factor AtHMGB2 as an enhancer of evoCAST-mediated integration activity in plants. These results establish CAST as a functional platform for programmable DNA insertion in plants and provide a foundation for developing targeted genome-engineering technologies for crop biotechnology.

## Introduction

Genome engineering has become a powerful tool for crop improvement, enabling precise modification of endogenous genes to accelerate breeding cycles and introduce agronomically valuable traits^1^. CRISPR-Cas nucleases and their derivatives now offer a suite of tools for targeted genomic modifications of varying scale^2^. At the small-edit level, Cas9- and Cas12-mediated indels efficiently knock out gene function^3–5^, while base editors enable precise single-nucleotide conversions without double-strand breaks (DSBs)^6,7^, and prime editors install defined substitutions, short insertions, and deletions with versatility^8–10^. More recently, hypercompact nucleases derived from IS200/IS605-family transposon-encoded proteins, such as TnpB and IscB, have been adapted for targeted mutagenesis in plants^11–14^.

Many agronomically important modifications, including transgene stacking, replacement of multi-kilobase regions, insertion of synthetic gene circuits, and molecular farming, require targeted integration of large DNA fragments. However, this capability remains a major unmet challenge in plant genome engineering^15,16^. Conventional *Agrobacterium*-mediated T-DNA transfer can deliver multi-kilobase fragments but integrates at random chromosomal positions^17^. Several strategies for targeted large-fragment insertion have been explored in plants. Homology-directed repair (HDR) induced by CRISPR-Cas DSBs can achieve precise knock-in, and multiple approaches have been developed to improve HDR rates, including geminivirus replicon-mediated donor amplification^18,19^, RNA virus-based delivery of repair templates^20^, tandem-repeat HDR designs^21^, and transcript-templated HDR^22^.

More recently, mobile-element enzymes have been repurposed for large DNA insertions in plant genome engineering. Recombinase-mediated approaches, such as PrimeRoot, which combines prime editing with site-specific recombination to insert fragments up to 11.1 kb in rice^23^, offer high precision but require the prior installation of recombinase recognition sites. DNA transposase-based systems use cut-and-paste transposition chemistry, which has been implemented by combining the rice Pong/mPing transposon system with Cas9 or Cas12a, allowing CRISPR-induced DSBs to guide mPing insertion at target loci in *Arabidopsis thaliana* and soybean^24^. A related but mechanistically distinct strategy uses an integration-defective piggyBac variant fused to Cas9 to tether donor DNA near Cas9-induced breaks and thereby enhance HDR-mediated knock-in in *Nicotiana benthamiana* and rice^25^. Lastly, engineered R2 retrotransposon-based platforms use a distinct mechanism in which an RNA intermediate is reverse-transcribed and inserted into conserved 25S rDNA loci, and have recently been demonstrated in *Arabidopsis thaliana, Nicotiana benthamiana*, and *Solanum lycopersicum*^26^, although their programmability still needs to be developed in plants.

CRISPR-associated transposases (CASTs) are another group of mobile elements that are promising for targeted insertion of large DNA payloads. They are naturally occurring bacterial systems that couple the RNA-guided target recognition of CRISPR effectors to the DNA integration activity of Tn7-like transposons, enabling programmable, DSB-free insertion of large DNA cargoes. Two major classes have been characterized: Type V-K systems, which use the single-effector Cas12k to direct transposition^27^, and Type I-F systems, which employ the multi-subunit Cascade complex (Cas8-Cas7-Cas6) for target recognition^28^. In Type I-F CASTs, QCascade, composed of Cascade subunits, crRNA, and TniQ, binds a crRNA-specified genomic target and recruits the AAA+ ATPase TnsC. Target-associated TnsC then recruits the heteromeric TnsA-TnsB transposase, which catalyzes transposon integration at a characteristic distance downstream of the crRNA-defined target site. TnsA and TnsB perform coordinated transposon-end processing and strand-transfer reactions that insert the donor transposon and generate 5-bp target-site duplications (TSD), a molecular hallmark of Tn7-family transposition^29–32^.

In bacteria, CAST systems achieve up to almost 100% integration frequency with selection^33,34^. Extending this capability to eukaryotes, Lampe et al. reconstituted Type I-F *Pseudoalteromonas* CAST (PseCAST) in human cells by engineering nuclear localization, optimizing QCascade function, screening CAST orthologs, and identifying the bacterial AAA+ unfoldase ClpX as an accessory factor that substantially enhances genomic integration efficiencies from ∼0.1% to ∼1%^35^. Witte et al. developed evoCAST by applying phage-assisted continuous evolution (PACE) to the transposase module and rationally engineering QCascade module. The evoCAST achieved 10-20% integration efficiencies in HEK293T cells, supported >10-kb cargo integration, and reduced the requirement for ClpX compared with wild-type PseCAST^36^.

Given the advantages of CASTs, including programmable target selection, independence from DSBs and host-repair pathways, and the capacity to carry multi-kilobase cargoes, establishing CAST activity in plants would open a new route for targeted DNA integration. Here, we demonstrated that Type I-F PseCAST can be engineered for targeted DNA integration in plant cells. We first established RNA-guided episomal integration in *A. thaliana* protoplasts, and then demonstrated targeted chromosomal integration at both transgenic and endogenous loci in *N. benthamiana*. Integration products retained canonical Tn7-family features, including 5-bp targetsite duplications and insertion positions centered approximately 49 bp downstream of the PAM. evoCAST further improved integration efficiency to 2.7%, representing a 6-fold increase over wild-type PseCAST, and supported crRNA-programmable targeting across six endogenous loci. A candidate accessory-factor screen identified AtHMGB2 as a chromatin-associated factor that enhances evoCAST activity, while targeted integration by a second Type I-F ortholog from *Vibrio cholerae* supports broader CAST portability in plant cells. Together, these results establish a foundation for CAST-mediated programmable cargo insertion in plant genomes.

## Results

### PseCAST components are reconstituted and expressed in plant cells

Type I-F CASTs require coordinated expression of a multi-component targeting and integration machinery. The PseCAST system can be functionally organized as two modules: i) a QCascade targeting module composed of Cas8, Cas7, Cas6, TniQ, and a programmable crRNA, and ii) an integration component composed of TnsA, TnsB, TnsC, and a mini-transposon donor carrying cargo flanked by the right-end (RE) and left-end (LE) sequences **(Fig. 1A)**.

**Figure 1.**
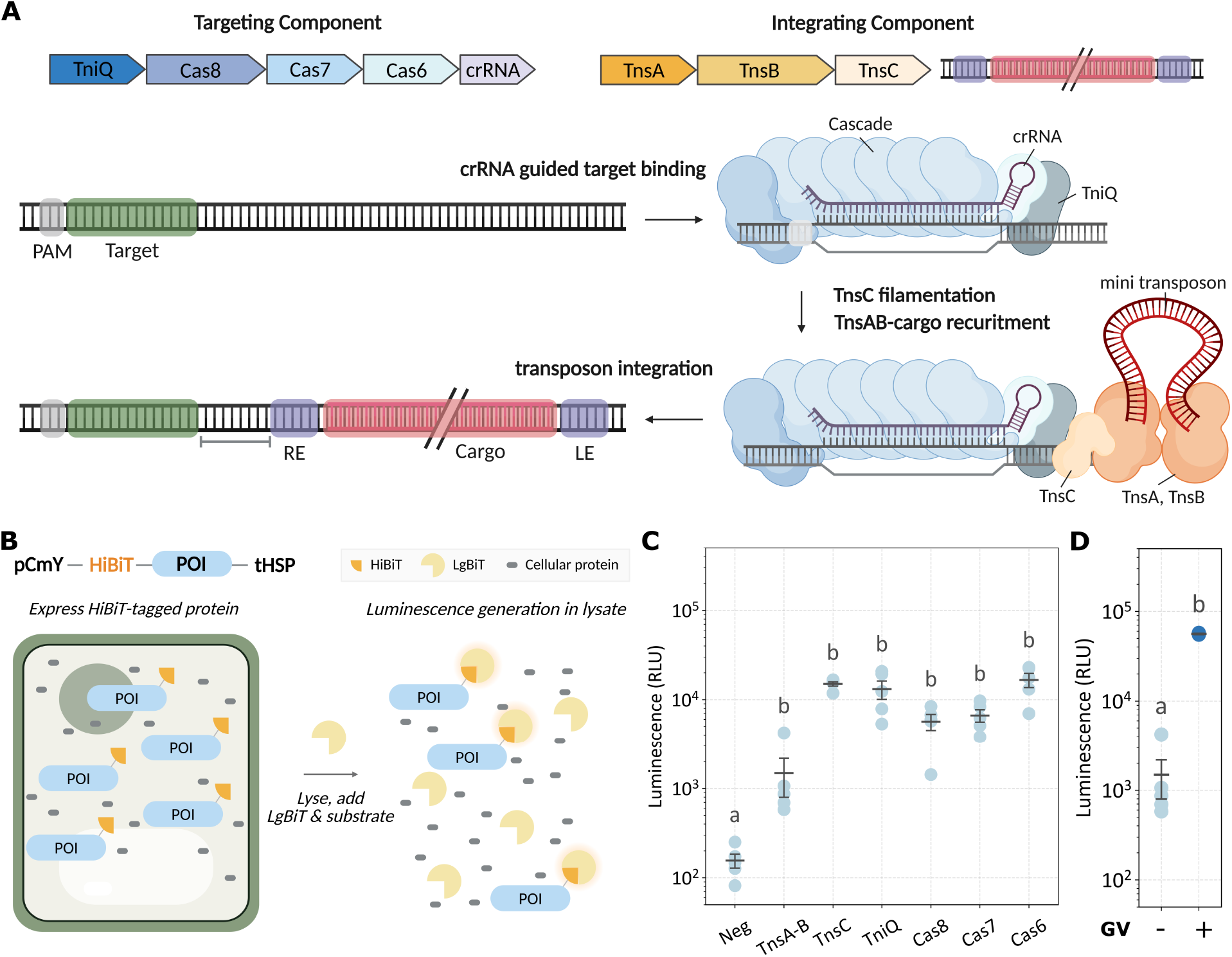
CAST system overview and expression of CAST components in plant cells. **(A)** Schematic of the Type I-F PseCAST system used for RNA-guided DNA integration. The targeting component consists of TniQ, Cascade subunits, and a crRNA that recognizes a genomic target adjacent to a PAM. Target-bound QCascade recruits TnsC, which promotes recruitment of the TnsA-TnsB transposase complex and insertion of a mini-transposon donor flanked by RE and LE sequences. Integration places the cargo downstream of the crRNA-defined target site and generates the characteristic Tn7-family insertion product. **(B)** Schematic of the HiBiT-based assay for detecting CAST component expression in plant cells. Each PseCAST protein was fused to an N-terminal HiBiT peptide tag and expressed under the pCmY promoter with the tHSP terminator. Following cellular fractionation, HiBiT complementation with LgBiT and substrate enables luminescence-based detection of each tagged protein. **(C)** Luminescence-based detection of individual HiBiT-tagged PseCAST components in plant nuclear fractions. **(D)** Geminiviral replicon (GV)-mediated enhancement of TnsA-TnsB detection in plant nuclear fractions. Bars represent mean ± S.E.M.; individual points indicate biological replicates (n=5). Statistical analysis was performed using one-way ANOVA with Tukey’s HSD post hoc test. Different letters indicate statistically significant differences (p < 0.05). pCmY: CmYLCV6 promoter, tHSP: HSP18.2 terminator, RLU: relative luminescence units.

We first tested whether the individual PseCAST proteins could be detected in plant nuclear fractions. Each PseCAST protein was fused to a strong bipartite nuclear localization signal and a HiBiT tag, and transiently expressed in *N. benthamiana* leaves via agroinfiltration using the pCmY promoter and the tHSP terminator. After cell lysis and nuclear enrichment, HiBiT complementation with exogenous LgBiT and substrate enabled luminescence-based detection of each tagged protein **(Fig. 1B)**. All tested CAST components produced luminescence signals above the negative background control. However, the TnsA-TnsB subunit produced a comparatively lower signal than the other CAST components **(Fig. 1C)**. To increase TnsA-TnsB expression, we used a geminiviral replicon (GV) to amplify the TnsA-TnsB expression cassette. The cassette was flanked by the long and short intergenic regions (LIR/SIR), and RepA was supplied to initiate replication^18^. This modification increased the TnsA-TnsB signal by approximately 40-fold relative to the non-GV configuration **(Fig. 1D)**. Together, these results demonstrate that PseCAST proteins can be expressed and detected in plant nuclear fractions and support the use of GV-based cassette amplification to increase expression of low-abundance CAST components.

### PseCAST mediates RNA-guided episomal integration in Arabidopsis thaliana protoplasts

To test whether PseCAST can achieve RNA-guided transposition in plant cells, we first performed an episomal integration assay in *A. thaliana* protoplasts. This episomal system allowed us to evaluate transposition activity without the confounding effects of chromosomal context. Because the combined size of all PseCAST expression cassettes exceeds the practical limits for delivery as a single plasmid through PEG transfection, the system was divided across two plasmids: pTarget-CAS (∼11 kb, encoding the Cascade subunits Cas8-Cas7-Cas6 and a crRNA targeting a site within the pTarget sequence, with mTurquoise as a transfection marker) and pDonor-TNP (∼14 kb, encoding TnsA-TnsB-TnsC-TniQ and the mini-transposon, with YPET as a transfection marker) **(Fig. 2A)**. crRNAs were expressed from U6 promoter-terminator cassettes, protein-coding subunits were expressed using pCmY promoter and tHSP terminator, and optimized nuclear localization sequences (NLS) from previous mammalian-cell CAST studies were incorporated accordingly^35^. We designed the mini-transposon cargo to carry a promoter-less mCherry flanked by RE and LE sequences, such that mCherry fluorescence is generated only when the cargo integrates downstream of the constitutive p35S promoter in pTarget **(Fig. 2A)**.

**Figure 2.**
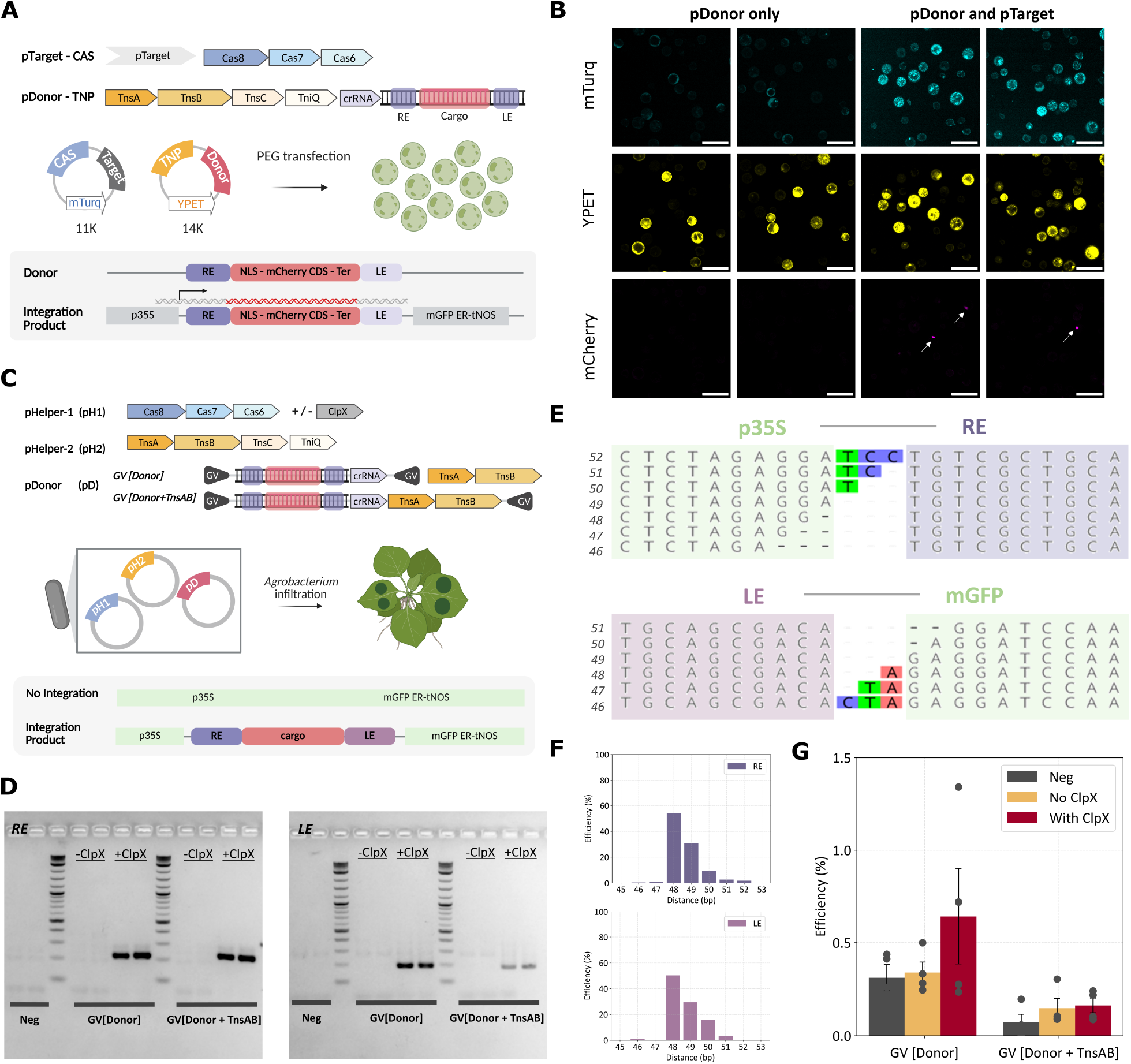
Wild-type PseCAST mediates RNA-guided episomal and chromosomal integrations. **(A)** Schematic of the episomal integration assay in *A. thaliana* protoplasts. CAST components were divided across two plasmids to accommodate the size of the system. **(B)** Representative fluorescence microscopy images of protoplasts receiving pDonor-TNP alone or both pDonor-TNP and pTarget-CAS. Arrows indicate mCherry-positive cells. **(C)** Schematic of the chromosomal integration assay in *N. benthamiana*. Three constructs were co-delivered by *Agrobacterium tumefaciens*. Two donor configurations were tested in parallel: GV-Donor, which contains the RE-cargo-LE mini-transposon and crRNA cassette, and GV-Donor+TnsAB, which additionally encodes supplemental TnsA-TnsB. **(D)** Nested PCR detection of chromosomal RE and LE junctions after delivery of wild-type PseCAST components. The negative control contains only the donor construct. **(E)** Representative NGS reads spanning the p35S-RE junction (top) and LE-mGFP junction (bottom). **(F)** Distribution of insertion distances at the RE (top) and LE (bottom) junctions. **(G)** Integration efficiency measured by droplet digital (ddPCR) for wild-type PseCAST using crRNA T1 across GV-Donor and GV-Donor+TnsAB configurations, with or without ClpX co-expression. Bars represent mean ± S.E.M.; individual points indicate biological replicates (n=4). Statistical analysis was performed using one-way ANOVA with Tukey’s HSD post hoc test. Different letters indicate statistically significant differences (p < 0.05). p35s: CaMV 35S promoter, mGFP ER: endoplasmic reticulum-targeted modified green fluorescent protein, tNOS: nopaline synthase terminator.

After protoplast transfection, fluorescence microscopy confirmed the presence of mTurquoise and YPET signals. Among cells receiving both pTarget-CAS and pDonor-TNP, a subset of them showed mCherry fluorescence **(Fig 2B)**. To confirm integration at the molecular level, we performed nested PCR using primer pairs spanning the right and left integration junctions **(Figure S1A)**. Expected junction products were detected for two independent crRNAs, T1 and T2, at both the RE and LE junctions, but only when pTarget-CAS and pDonor-TNP were co-delivered **(Figure S1B, C)**. A time-course experiment further revealed integration products detectable as early as 12 h after transfection, with band intensity increasing at 48 h **(Figure S1D)**. These results show that PseCAST mediates RNA-programmable episomal transposition in *A. thaliana* protoplasts.

### PseCAST integrates into a transgenic chromosomal locus in N. benthamiana

We next tested whether PseCAST could integrate cargo into a chromosomal target. We used the transgenic 16C *N. benthamiana* line as an initial target because it carries a p35S-mGFP-tNOS transgene that could be targeted using the same crRNAs validated in the episomal assay^37^. For *Agrobacterium*-mediated infiltration, CAST components were divided across two Helper and one Donor constructs. The first helper construct encoded Cas8, Cas7, and Cas6, with or without ClpX. The second helper construct encoded the transposition-associated components TnsA-TnsB, TnsC, and TniQ. The donor construct contained a 200-bp synthetic cargo sequence flanked by RE and LE, and incorporated geminiviral replicon (GV) to increase free donor DNA copy number **(Fig. 2C)**. Two donor configurations were tested in parallel: GV [Donor], which carried the RE-mCherry-LE transposon together with a crRNA cassette; and GV [Donor+TnsAB], which additionally encoded TnsA-TnsB expression cassette to compensate for the comparatively lower TnsA-TnsB signal observed in the HiBiT assay **(Fig. 1C, Fig. 2C)**.

Three constructs were co-infiltrated into *N. benthamiana* leaves, and genomic DNA was extracted 7 days after infiltration. Nested PCR across the RE and LE junctions detected chromosomal integration products only in leaves infiltrated with the ClpX-containing helper plasmids **(Fig. 2D)**. Amplicon sequencing of the junction products confirmed hallmark Tn7-family integration features, including 5-bp TSDs and insertion positions concentrated around 48-49 bp downstream of the crRNA-directed target site **(Fig. 2E, F)**. We performed probe-based droplet digital PCR (ddPCR) targeting the RE integration junction and single-copy *NbRDR1* reference gene, with integration frequency calculated by normalizing the integrated junction copy number to the reference gene copy number. PseCAST produced detectable but less than 1% integration efficiencies under these conditions **(Fig. 2G, Figure S2A)**. We further tested another crRNA, T2, targeting the p35S-mGFP-tNOS transgene. Amplicon sequencing indicated the expected junctions and insertion-distance profiles, confirming canonical integration at the second target site **(Figure S2B, C)**. Probe-based quantitative PCR (qPCR) and ddPCR yielded variable, low-level signals, reflecting the limited activity of wild-type PseCAST in plant systems **(Figure S2D, E)**.

We also compared Cascade expression architectures, including monocistronic and 2A-linked polycistronic configurations **(Figure S3A)**^38^. All tested designs supported detectable integration by nested PCR, and the monocistronic pCmY-tHSP configuration showed the strongest apparent activity, although differences among architectures were not statistically significant at this low level of integration efficiency **(Figure S3B, C)**.

### Additional CAST orthologs show low activity in plants

We next asked whether additional CAST systems could function in plant cells. We first tested VchCAST from *Vibrio cholerae* Tn6677 using an analogous chromosomal integration assay targeting the 16C *N. benthamiana*^34,35^. HiBiT-based detection confirmed that all VchCAST proteins could be expressed and recovered from plant nuclear fractions, although TnsA-TnsB again produced comparatively lower signals and could be increased with the use of GV **(Figure S4A, B)**. Nested PCR across the RE integration junction detected chromosomal integration products in both the GV [Donor] and GV [Donor+TnsAB] configurations when ClpX was included. ddPCR further confirmed low but reproducible insertion events compared to the negative controls without Helper constructs **(Figure S4C, D)**.

We also evaluated Homing Endonuclease-assisted Large-sequence Integrating CAST complex (HELIX), an engineered Type V-K CAST architecture derived from the *Scytonema hofmanni* ShCAST system^27,47^. The HELIX design incorporates an engineered nicking module, LAGLIDADG homing endonuclease (LHE) from *Aspergillus nidulans* (I-Anil), into the transposase machinery, thereby promoting more complete donor excision^48,49^. For the episomal integration assay, HELIX components were divided across two plasmids: pDonor-CAS, encoding Cas12k, the crRNA, and the mini-transposon cargo, and pTarget-TNP, encoding the target site and HELIX-derived transposition machinery. Four transposition machinery configurations were tested, differing in NLS and I-Anil placement and intron usage **(Figure S4E)**. Nested PCR across the RE and LE integration junctions failed to detect the expected amplicons across all tested configurations, suggesting that additional optimization is required to establish the Type V-K system in plants **(Figure S4F)**.

### EvoCAST improves integration efficiency in protoplasts and leaves

To improve CAST activity in mammalian cells, Witte et al. developed evoCAST, an optimized PseCAST variant that combines PACE-evolved transposase components with rationally engineered QCascade components^36^. The PACE-derived substitutions are distributed across TnsA, TnsB, and TnsC, with the TnsB mutations likely playing a major role because they map to predicted interfaces involved in transposon-end binding, TnsB-TnsB assembly, and TnsC-mediated recruitment. The rationally engineered QCascade component introduced structure-guided PAM-contact mutations in Cas8 together with DNA-contact mutations in Cas7, and added bipartite NLSs to Cas8, Cas6, and TniQ to strengthen nuclear import **(Figure S5A)**. We first validated evoCAST activity in the episomal protoplast assay. Similar to wild-type PseCAST, fluorescence microscopy detected mCherry expression in a few co-transfected protoplasts **(Fig. 3A)**. Nested PCR using primer pairs spanning the right and left integration junctions produced bands of the expected sizes for both crRNA T1 and crRNA T2 **(Figure S5B, C)**, providing molecular evidence that evoCAST preserves crRNA-directed integration specificity in plant cells.

**Figure 3.**
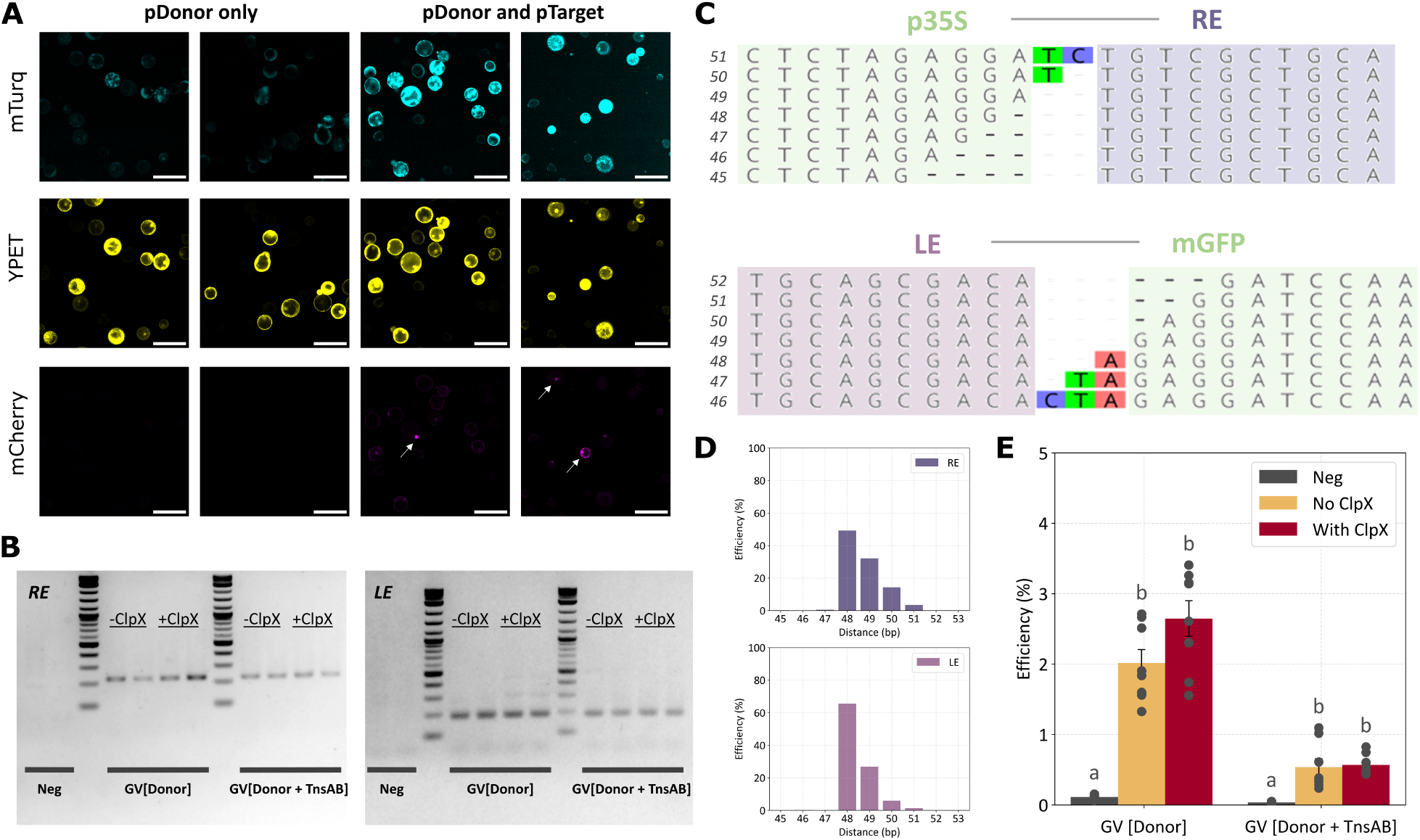
EvoCAST improves integration efficiency in plants. **(A)** Representative fluorescence microscopy images of protoplasts receiving pDonor-TNP alone or both pDonor-TNP and pTarget-CAS. mTurquoise and YPET fluorescence indicate plasmid delivery. Arrows indicate mCherry-positive cells. **(B)** Nested PCR detection of chromosomal RE and LE junctions after *Agrobacterium*-mediated delivery of evoCAST components to *N. benthamiana* leaves. The negative control contains only the donor construct. **(C)** Representative NGS reads spanning the p35S-RE junction (top) and LE-mGFP junction (bottom) from panel B experiments. **(D)** Distribution of insertion distances at the RE (top) and LE (bottom) junctions. **(E)** Integration efficiency measured by droplet digital PCR (ddPCR) for evoCAST using crRNA T1 across GV [Donor] and GV [Donor+TnsAB] configurations, with or without ClpX co-expression. Bars represent mean ± S.E.M.; individual points indicate biological replicates (n=8). Statistical analysis was performed using one-way ANOVA with Tukey’s HSD post hoc test. Different letters indicate statistically significant differences (p < 0.05). GV: geminiviral replicon.

We next evaluated evoCAST at the chromosomal p35S-mGFP-tNOS target in the 16C *N. benthamiana*. Nested PCR detected RE and LE integration junction products in both GV [Donor] and GV [Donor+TnsAB], with products observed in both the absence and presence of ClpX **(Fig. 3B)**. This is consistent with prior mammalian-cell results showing that evoCAST largely reduced the requirement for ClpX^36^. Amplicon sequencing confirmed that evoCAST retained canonical transposition signatures, including 5-bp TSDs at both integration ends and an insertion-distance distribution centered around 48-49 bp downstream of the target site **(Fig. 3C, D)**.

ddPCR quantification showed that evoCAST significantly increased the efficiency of chromosomal integration relative to wild-type PseCAST. In the GV [Donor] configuration, evoCAST achieved 2.0 and 2.7% integration without and with ClpX, representing 5.9-fold and 4.1-fold improvements, respectively, over wild-type PseCAST under comparable delivery conditions **(Fig. 3E)**. The GV [Donor] configuration produced higher overall integration efficiency than GV [Donor+TnsAB] configuration, suggesting that additional TnsA-TnsB expression was not helpful for evoCAST activity. Similar amplicon sequencing, qPCR, and ddPCR results were obtained with crRNA T2 **(Figure S6)**. Based on these results, the GV [Donor] configuration with ClpX co-expression was used for subsequent experiments. These results demonstrate that evoCAST improves CAST integration activity and reduces ClpX dependence, while preserving canonical RNA-guided transposition signatures in the plant cellular environment.

### EvoCAST enables cis-regulatory element insertion at a synthetic landing pad

Targeted insertion of cis-regulatory elements provides a potential strategy for tuning gene expression and engineering agronomic traits without modifying the underlying protein-coding sequence^39^. We asked whether a promoter could serve as a functional evoCAST cargo and be installed at a predefined site. To test this concept, we designed a synthetic landing pad in which YPET and DsRed coding sequences flanked an intergenic target region and faced in opposite directions. The donor carries a UBIQUITIN10 promoter (pUBQ10) between the RE and LE sequences, while the crRNA targets the intergenic region. Depending on the integration orientation, the inserted promoter would activate either YPET or DsRed expression **(Fig. 4A)**.

**Figure 4.**
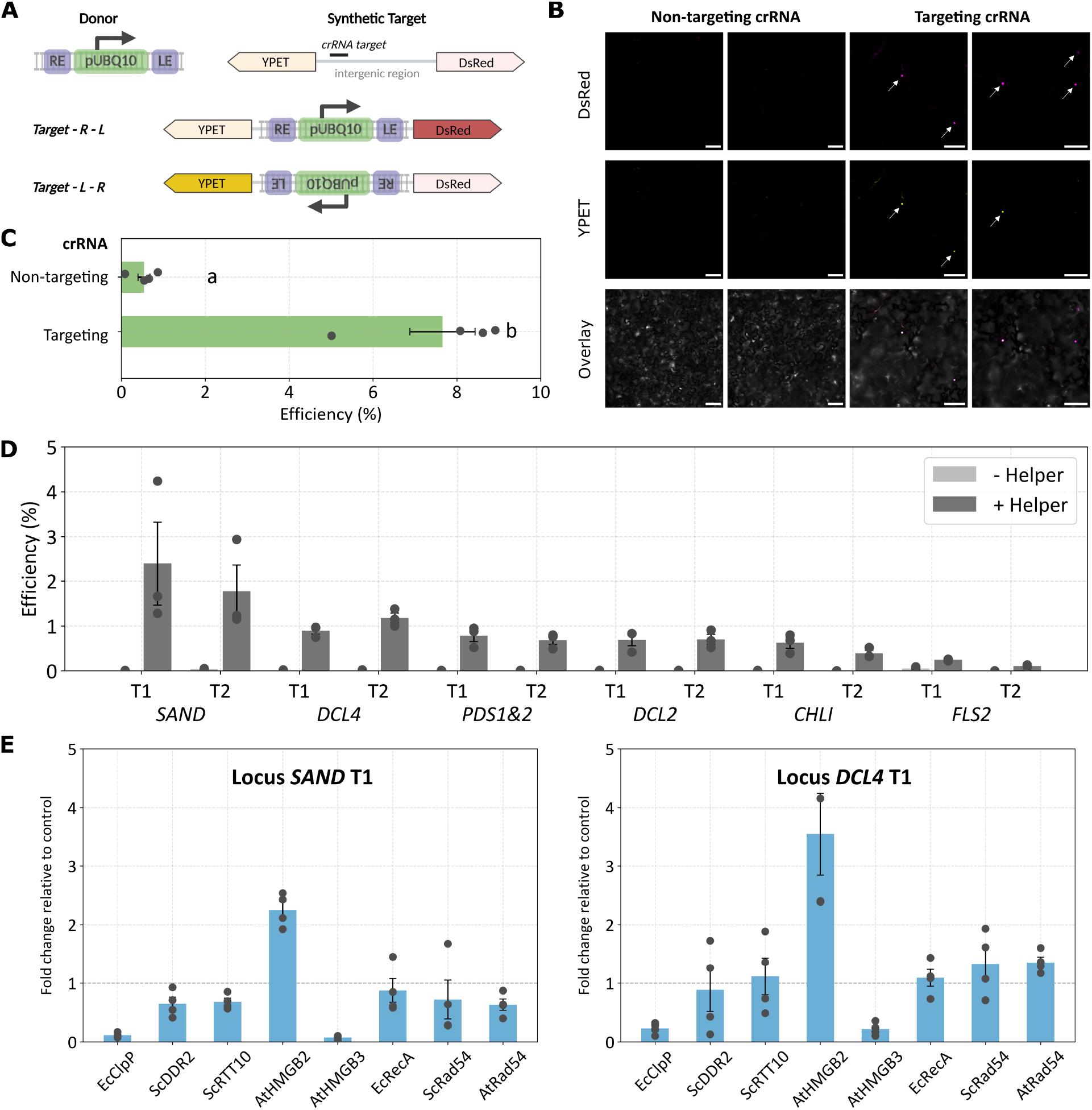
EvoCAST achieves integration at a synthetic landing pad and endogenous loci. **(A)** Schematic of the promoter integration assay. Promoter-less YPET and DsRed coding sequences flank an intergenic region containing the crRNA target and face in opposite directions. The donor carries the pUBQ10 promoter. Integration in the R-L orientation activates DsRed expression, whereas integration in the L-R orientation activates YPET expression. **(B)** Representative confocal fluorescence images of *N. benthamiana* leaves following delivery of evoCAST with either a non-targeting or a targeting crRNA. Arrows indicate DsRed- and YPET-positive cells. **(C)** Quantification of integration efficiency for the preferred R-L orientation by ddPCR. Bars represent mean ± S.E.M.; individual points indicate biological replicates (n=4). Statistical analysis was performed using one-way ANOVA with Tukey’s HSD post hoc test. Different letters indicate statistically significant differences (p < 0.05). **(D)** Quantification of evoCAST integration efficiency across six endogenous *N. benthamiana* loci (*SAND, DCL4, PDS1&2, DCL2, CHLI*, and *FLS2*) by qPCR. Each locus was targeted with two crRNAs (T1 and T2). Bars represent mean ± S.E.M.; individual points indicate biological replicates (n=3). **(E)** Fold change in evoCAST integration efficiency upon co-expression of candidate cofactors, measured by qPCR at *SAND* T1 (left) and *DCL4* T1 (right). Bars represent mean ± S.E.M.; individual points indicate biological replicates (n=4). pUBQ10: UBIQUITIN10 promoter.

Following *Agrobacterium*-mediated infiltration of evoCAST Helper and Donor constructs with the synthetic landing pad into *N. benthamiana* leaves, the targeting crRNA generated both DsRed- and YPET-positive cells, whereas no fluorescent cells were observed with the non-targeting crRNA control **(Fig. 4B)**. Fluorescent images revealed a preference for R-L orientation, in which the donor RE is positioned proximal to the target site and LE is positioned distal to the target site, as shown by greater number of cells expressing DsRed compared to YPET **(Fig. 4B)**. Quantification of the preferred target-R-L orientation via ddPCR showed an integration efficiency of approximately 8% with the targeting crRNA, and less than 1% off-target integration with the non-targeting crRNA **(Fig. 4C)**.

### EvoCAST mediates crRNA-programmable integration at endogenous N. benthamiana loci

Having established evoCAST activity at both a transgenic target and a synthetic landing pad, we next asked whether the system could be targeted to endogenous chromosomal loci. We designed two crRNAs for each of six *N. benthamiana* loci: *SAND family protein (SAND), DICER-LIKE 4 (DCL4), PHYTOENE DESATURASE 1 and 2 (PDS1 and PDS2), DICER-LIKE 2 (DCL2), MAGNESIUM CHELATASE SUBUNIT I (CHLI)*, and *FLAGELLIN-SENSITIVE 2 (FLS2)* **(Fig. 4D)**. Each crRNA was incorporated into the same GV [Donor] delivery architecture, and integration was quantified by probe-based qPCR. evoCAST-mediated integration was detected above the no-helper control at all six endogenous loci, indicating that the system can be retargeted by changing the crRNA sequence alone. Integration efficiency varied across loci and crRNAs. *SAND* showed the highest activity in this panel, with efficiencies reaching 2.1%, whereas *FLS2* showed the lowest activity, remaining below 0.18% under the tested conditions **(Fig. 4D)**.

These results demonstrate that evoCAST activity is not restricted to a synthetic landing pad and can be programmed to insert cargo at diverse endogenous positions in the *N. benthamiana* genome.

### Chromatin-associated cofactors modulate evoCAST integration efficiency

The locus-dependent activity observed across endogenous targets suggested that local chromosomal context may influence CAST integration. To identify factors that could improve evoCAST performance, we screened candidate accessory proteins associated with protein turnover (EcClpP)^35,40^, stress and transposition-related responses (ScDDR2 and ScRTT10)^41^, chromatin architecture (AtHMGB2 and AtHMGB3)^42,43^, and recombination-associated DNA repair or chromatin remodeling (EcRecA, ScRad54, and AtRad54)^44–46^ **(Fig. 4E)**. Integration was quantified at two endogenous targets, *SAND* T1 and *DCL4* T1, as fold change relative to the control where no cofactor is included. Among the tested candidates, AtHMGB2 produced the strongest enhancement at both loci, increasing integration by 2.3-fold at *SAND* T1 and 3.5-fold at *DCL4* T1. By contrast, EcClpP and AtHMGB3 reduced integration relative to the control under the tested conditions **(Fig. 4E)**.

## Discussion

Targeted DNA integration remains a major unmet challenge in precision plant genome engineering. Here, we demonstrate that Type I-F CAST can be reconstituted in plants to achieve RNA-guided DNA integration. The first challenge was the coordinated expression of the multi-component transposition machinery. Productive activity requires nuclear accumulation of QCascade, TniQ, TnsC, TnsA-TnsB, crRNA, and donor DNA, making both component abundance and stoichiometry likely determinants of integration efficiency. Our HiBiT assay showed detectable expression of all tested PseCAST components in plant nuclear fractions, but TnsA/TnsB produced a lower signal **(Fig. 1C)**. Geminiviral replicon amplification substantially increased the TnsA-TnsB HiBiT signal; however, in chromosomal integration assays, the GV [Donor+TnsAB] configuration was less efficient than the GV [Donor] configuration alone **(Fig. 1D, Fig. 2G, and Fig 3E)**. One likely explanation is that adding the TnsA-TnsB cassette increased the size of the GV replicon, which can reduce replicon copy number and donor availability^50^. This result indicates that increasing the abundance of a limiting component does not necessarily improve integration and may instead alter the stoichiometric balance. This finding is consistent with mammalian cell PseCAST studies showing that eukaryotic reconstitution requires optimization of expression architecture, nuclear localization signal placement, plasmid stoichiometry, and relative component abundance^35^. Future optimization should therefore focus not only on stronger promoters, but also on balancing the targeting, transposition, and donor modules to favor productive complex assembly.

The transition from bacterial CAST activity to eukaryotic genome integration highlights both host-specific constraints and conserved mechanistic bottlenecks. In mammalian cells, Lampe et al. identified post-transposition complex resolution as a key genomic-integration bottleneck, showing that bacterial ClpX enhanced genomic integration by approximately two orders of magnitude while leaving episomal integration largely unaffected^35^. This finding was motivated by bacterial transposition biology, where stable post-transposition complexes have been described for Tn7 and Mu, and ClpX is known to disassemble the Mu transposase complex to promote downstream gap repair^51,52^. Consistent with this model, ClpX co-expression enabled the detection of wild-type PseCAST chromosomal integration in *N. benthamiana* **(Fig. 2D)**.

evoCAST improves the intrinsic activity of the CAST machinery through PACE evolution and rational engineering. In plants, evoCAST increased integration efficiency by approximately 4-6-fold over wild-type PseCAST and reduced ClpX dependence. This result suggests that some rate-limiting molecular interactions are conserved across bacterial, mammalian, and plant contexts. However, the absolute efficiencies observed in plants, approximately 2.5% **(Fig. 3E)**, remained lower than the 10%-20% efficiencies reported for optimized evoCAST in mammalian cells. We hypothesize that this difference may reflect plant-specific barriers, including constraints imposed by *Agrobacterium*-mediated delivery, geminiviral replicon size and copy number, the absence of edited-cell enrichment, and plant chromatin state.

Chromatin context shapes the activity of genome-engineering systems in eukaryotic cells. Across the six endogenous loci tested, evoCAST integration efficiency varied by approximately an order of magnitude, ranging from 2%-2.5% at the highest-activity loci to 0.3% at the lowest-activity locus **(Fig. 4D)**. This target-dependent activity is consistent with mammalian evoCAST results, where Witte et al. reported a positive correlation between integration efficiency and chromatin accessibility measured by ATAC-seq across 14 genomic sites^53^. Chromatin effects have also been observed in plant genome editing: in an *A. thaliana* reporter system, identical Cas9 target sequences placed in different chromatin environments showed up to ∼250-fold differences in mutagenesis efficiency, with accessible euchromatic features generally associated with higher editing and heterochromatic features, DNA methylation, and low accessibility associated with reduced editing^54^. Although CAST integration differs mechanistically from Cas9 cleavage, both systems require a large RNA-guided protein complex to access chromosomal DNA. Thus, local accessibility, nucleosome occupancy, transcriptional activity, and DNA methylation state may similarly affect QCascade binding, TnsC recruitment, donor capture, or post-integration repair. By comparison, in the promoter insertion assay at the synthetic landing pad, integration efficiency reached approximately 8% **(Fig. 4C)**. Because the synthetic target was transiently delivered by *Agrobacterium*, the presence of multiple T-DNA copies or extrachromosomal T-DNA molecules may have provided additional integration substrates, contributing to the higher measured efficiency^55,56^.

This target-site dependence also highlights a potential application of plant CAST for safe harbor discovery. Ideal plant safe harbors should support stable transgene expression, reproducible performance, and minimal growth effects, yet experimentally validated loci remain limited in many crop species^23^. Compared with HDR-based knock-in, CAST could make candidate-site testing more scalable because loci can be retargeted by changing only the crRNA while keeping the donor and transposition machinery constant, but integration efficiencies will likely need to be improved for this application.

These considerations motivated us to screen host-cell cofactors that could improve one or more of these steps. Among the candidates tested, AtHMGB2 produced the strongest enhancement at both *SAND* T1 and *DCL4* T1, whereas the related AtHMGB3 instead reduced integration efficiency **(Fig. 4E)**. HMGB proteins are non-histone chromatin architectural proteins that bind and bend DNA and function as versatile co-regulators^57^. This function provides a plausible mechanistic link to CAST integration, which requires coordinated target recognition, TnsC recruitment, transposon-end engagement, strand transfer, and host-mediated repair. Prior work showed that mammalian HMGB1 acts as a cellular cofactor for Sleeping Beauty transposition by promoting transposase-transposon synaptic complex formation^42^. Although AtHMGB2 and AtHMGB3 share approximately 87% sequence similarity, their contrasting effects indicate that CAST enhancement effect is paralog-specific^57^. Plant HMGB proteins can differ in nuclear distribution, chromatin-binding dynamics, acidic-tail regulation, and protein-interaction behavior, any of which could influence CAST activity.

Several additional features of plant evoCAST remain to be characterized. First, our synthetic promoter integration assay produced both DsRed- and YPET-positive cells **(Fig. 4B)**, whereas Type I-F CAST systems generally exhibit a strong preference for the right-to-left integration orientation, and evoCAST retained this orientation bias^31,53^. The integration-orientation preference should be examined further in plant cells. Second, in human cells, transient evoCAST delivery produced QCascade-independent off-target integration by highly active TnsA-TnsB-TnsC complexes, whereas off-target integration was not detected after selection for on-target AAVS1-edited cells^53^. Specificity analysis is more challenging in plants because *Agrobacterium* delivery and geminiviral replicon amplification can leave abundant unintegrated donor DNA, increasing background in genome-wide assays. In addition, although our junction sequencing confirmed canonical insertion positions and target-site duplications, it did not quantify insertion orientation, simple insertion frequency, or cointegrate formation. Third, future work should characterize the heritability of the CAST-mediated insertions in plants.

Finally, our comparison of alternative CAST architectures suggests that plant-cell compatibility is modular and system-dependent. VchCAST generated target-specific integration products in *N. benthamiana*, indicating that plant CAST activity is not unique to the Pseudoalteromonas system. However, its lower efficiency suggests that differences among Type I-F orthologs in expression, QCascade targeting, TnsC recruitment, transposase activity, or post-integration processing are compatible with the plant nuclear environment. By contrast, the tested Type V-K HELIX configurations did not produce detectable episomal integration products, indicating that successful transfer of one CAST class does not guarantee portability of another. These results support a modular engineering strategy for plant genome editors: rather than testing whole systems as indivisible units, future efforts should optimize separable modules, including delivery, nuclear localization, target recognition, effector recruitment, catalytic activity, donor architecture, chromatin access, and host-factor compatibility. Such a framework may accelerate not only CAST development but also the adaptation of other multi-component genome-editing platforms to plants.

## Methods

### Plant materials and growth conditions

*Nicotiana benthamiana* and *Arabidopsis thaliana* plants were grown in a Conviron controlled-environment chamber set to 22 °C during the day and 20 °C during the night, with a 16 h light /8 h dark photoperiod and 55% relative humidity. 3-week-old *N. benthamiana* plants were used for leaf infiltrations for transient experiments. 4-weeks-old *A. thaliana* plant leaves were harvested for protoplast isolation.

### Construct design and cloning

All coding sequences were codon-optimized for expression in *N. benthamiana* using the GenSmart™ Codon Optimization tool. Plasmids were assembled with the Loop Assembly method using BsaI- and SapI-mediated Golden Gate reactions (NEB catalog numbers R0569S and E1601L) and cloned into pCAMBIA backbones. All plasmids were propagated in NEB Turbo *Escherichia coli* competent cells (NEB catalog no. C2984I) and sequence-verified by nanopore sequencing (Plasmidsaurus). Sequences and plasmids generated in this work are listed in **Supplementary Table 1 and 2**.

### HiBiT-based detection of CAST component expression

Protein coding sequences were fused to an N-terminal HiBiT peptide tag for higher throughput luminescence detection using the Nano-Glo® HiBiT Lytic Detection System. At 3 days post infiltration, three infiltrated *Nicotiana benthamiana* leaves were harvested, flash-frozen in liquid nitrogen, and ground to a fine powder before extraction in lysis buffer: 20 mM Tris-HCl pH 7.4 (ThermoFisher Scientific catalog no. 15567027), 25 % glycerol (Sigma-Aldrich catalog no. G5516), 20 mM KCl (Sigma-Aldrich catalog no. P5405), 2 mM EDTA (Sigma-Aldrich catalog no. E9884), 2.5 mM MgCl_2_(Sigma-Aldrich catalog no. M2393), 250 mM sucrose (ThermoFisher scientific catalog no. J65148.A1), 0.1 % PMSF (ThermoFisher Scientific catalog no. 36978). After centrifugation at 1500 × g for 10 min at 4 °C to pellet nuclei, the pellet was washed up to 5 times with wash buffer: 20 mM Tris-HCl pH 7.4 (ThermoFisher Scientific catalog no. 15567027), 25 % glycerol (Sigma-Aldrich catalog no. G5516), 2.5 mM MgCl_2_(Sigma-Aldrich catalog no. M2393), 0.2 % Triton X-100 (Sigma-Aldrich catalog no. S5886) and lysed in high-salt buffer (20 mM HEPES-KOH pH 7.9 (Sigma-Aldrich catalog no. H3375), 2.5 mM MgCl_2_(Sigma-Aldrich catalog no. M2393), 100 mM NaCl (Sigma-Aldrich catalog no. S5886), 20 % glycerol (Sigma-Aldrich catalog no. G5516), 0.2 mM EDTA (Sigma-Aldrich catalog no. E9884), 0.5 mM DTT (GOLDBIO catalog no. DTT), one cOmplete Mini EDTA-free protease-inhibitor tablet (Sigma-Aldrich catalog no. 11836170001) per 10 mL. For HiBiT detection, 30 µL of nuclear lysate was combined with 30 µL Nano-Glo HiBiT Lytic reagent (Promega catalog no. N3030) containing LgBiT and furimazine, and luminescence was recorded on a Tecan Spark plate reader with a 1000 millisecond integration time.

### A. thaliana protoplast isolation and PEG-mediated transfection

Mesophyll protoplasts were isolated from fully expanded leaves of *A. thaliana* following Yoo et al. with minor modifications^58^. On the day of the experiment, a fresh enzyme solution (20 mL) containing 20 mM MES pH 5.7 (Sigma-Aldrich catalog no. M2933), 0.4 M mannitol (Millipore Sigma catalog no. 443907), 20 mM KCl (Sigma-Aldrich catalog no. P5405), 1.5% (w/v) cellulase onozuka R-10 (Yakult Pharmaceutical IND. CO., LTD) and 0.4% (w/v) macerozyme R-10 (Yakult Pharmaceutical IND. CO., LTD) was prepared; the MES–mannitol–KCl base was pre-heated to 70°C for 4 min before the enzymes were dissolved. After cooling, 10 mM CaCl_2_(Sigma-Aldrich catalog no. C7902) and 0.1% (w/v) BSA (Fisher bioreagents catalog no. BP9703) were added, and the solution was passed through a 0.45 µm syringe filter into a Petri dish. *The leaves* were incubated in this solution without agitation for 3 h at room temperature in the dark, with gentle swirling once per hour. The digest was diluted 1:1 with W5 solution: 154 mM NaCl (Sigma-Aldrich catalog no. S5886), 125 mM CaCl_2_(Sigma-Aldrich catalog no. C7902), 5 mM KCl (Sigma-Aldrich catalog no. P5405), 2 mM MES pH 5.7 (Sigma-Aldrich catalog no. M2933), passed through a 70 µm nylon mesh (Fisherbrand catalog no. 22363548), and the flow-through was centrifuged at 200 × g for 2 min. The pellet was gently resuspended in 5 mL W5, chilled on ice for 30 min, then the supernatant was removed, and the protoplasts were adjusted to 5 × 105 cells mL^−1^ in MMG solution: 0.4 M mannitol (Millipore Sigma catalog no. 443907), 15 mM MgCl_2_(Sigma-Aldrich catalog no. M2393), 4 mM MES pH 5.7 (Sigma-Aldrich catalog no. M2933.

For PEG-mediated transfection, 40% PEG 4000 solution (Sigma-Aldrich catalog no. 81240) was freshly prepared: 4 g PEG 4000, 3 mL H_2_O, 2.5 mL 0.8 M mannitol (Millipore Sigma catalog no. 443907), 1 mL 1 M CaCl_2_(Sigma-Aldrich catalog no. C7902). In round-bottom Eppendorf tubes, ∼20 µg of plasmid DNA (≤ 20 µL) was combined with 100 µL of protoplast suspension, mixed gently, and 120 µL of PEG solution was added. After 15 min at room temperature, 1 mL W5 was added, and the mixture was centrifuged (50 × g, 2 min); the wash was repeated three times. The pellet was resuspended in WI solution: 0.5 M mannitol (Millipore Sigma catalog no. 443907), 20 mM KCl (Sigma-Aldrich catalog no. P5405), 4 mM MES pH 5.7 (Sigma-Aldrich catalog no. M2933), transferred to BSA-blocked well plates (0.1 % BSA (Fisher bioreagents catalog no. BP9703), and incubated at room temperature in the dark for 24-72 h before downstream analysis.

### Episomal integration assay and fluorescence detection in A. thaliana protoplasts

Isolated *A. thaliana* protoplasts were co-transfected with 2 pmol each of the pTarget-CAS and pDonor-TNP constructs per 100 µL of protoplast suspension at concentration of 5 × 10^5^ cells mL^−1^. Following transfection, 40 µL of the 100 µL protoplast suspension was transferred to a 96-well plate for fluorescence analysis. At 72 h post-transfection, protoplasts were imaged using a Leica STELLARIS 8 FALCON laser-scanning confocal microscope equipped with an HC PL APO 20×/0.75 NA water-immersion objective. Fluorescent proteins were excited at 434 nm for mTurquoise, 517 nm for YPET, and 587 nm for mCherry. Images were acquired at 20× magnification using identical acquisition settings within each experiment to allow quantitative comparison across samples.

### Agrobacterium-mediated leaf infiltration and chromosomal integration assay in N. benthamiana

*Agrobacterium tumefaciens* GV3101 strains harboring circuit constructs were cultured overnight at 30 °C with shaking at 200 rpm in 2×YT medium (ThermoFisher 22712020) supplemented with 10 µg mL^−1^ rifampicin (Sigma-Aldrich R3501), 20 µg mL^−1^ gentamicin (Sigma-Aldrich G1264), 50 µg mL^−1^ tetracycline (Sigma-Aldrich T7660), and 50 µg mL^−1^ kanamycin (Sigma-Aldrich K1637). Cultures were pelleted by centrifugation at 4,000 g for 5 min and resuspended in infiltration buffer (10 mM MES, pH 5.7; 10 mM MgCl_2_; 200 µM acetosyringone) to a final OD_600_of 0.6. Resuspended cultures were incubated at room temperature with shaking at 120 rpm for 4 h to induce virulence gene expression prior to infiltration.

For chromosomal integration assays, two helper constructs and one donor construct were co-delivered by mixing the corresponding *Agrobacterium* suspensions at equal volume ratios. Donor configurations with and without ClpX were included as indicated. Mixed *Agrobacterium* suspensions were infiltrated into the abaxial surface of fully expanded leaves of 3-week-old *N. benthamiana* plants using a needleless syringe. Biological replicates represent independent plants of the same age with leaves at comparable developmental stages. Leaf tissue was harvested 6 days after infiltration, and genomic DNA was extracted using the DNeasy Plant Mini Kit according to the manufacturer’s protocol.

### Nested PCR and Sanger sequencing

Genomic DNA was extracted using the DNeasy Plant Mini Kit (Qiagen, catalog no. 69106) according to the manufacturer’s protocol. For the first-round outer PCR, 10 µL reactions were assembled using 5 µL Phusion High-Fidelity PCR Master Mix (NEB, catalog no. M0531S), 0.5 µL each of forward and reverse primers, 0.3 µL DMSO, and 20 ng gDNA, with nuclease-free water added to a final volume of 10 µL. Primers are listed in **Supplementary Table 3**. Cycling condition was as follows: 98 °C for 30 s; 30 cycles of 98 °C for 10 s, 65 °C for 20 s, and 72 °C for 15 s; followed by a final extension at 72 °C for 2 min. The outer PCR product was diluted 50-fold and used as the template for the second-round inner PCR. Inner PCR reactions were assembled in 10 µL volumes containing 5 µL Phusion High-Fidelity PCR Master Mix, 0.5 µL each of forward and reverse primers, 0.3 µL DMSO, and 1 µL diluted outer PCR product, with nuclease-free water added to 10 µL. Cycling condition was as follows: 98 °C for 30 s; 30 cycles of 98 °C for 10 s, 68 °C for 20 s, and 72 °C for 10 s; followed by a final extension at 72 °C for 2 min. Final amplicons were verified by electrophoresis on a 2% agarose gel, purified using the QIAquick Gel Extraction Kit (Qiagen, catalog no. 28704), and submitted for Sanger sequencing.

### Next-generation sequencing of CAST integration junctions

CAST-mediated integration junction amplicons were resolved by agarose gel electrophoresis, gel-purified using the QIAquick Gel Extraction Kit (Qiagen, catalog no. 28704), and eluted in 35 µL nuclease-free water. DNA concentration was measured using NanoDrop 2000 spectrophotometer, and each amplicon was adjusted to 50 ng µL^−1^. Purified amplicons were sealed in Eppendorf microcentrifuge tubes and shipped overnight to the Massachusetts General Hospital CCIB DNA Core for Illumina sequencing. Sequencing reads were demultiplexed and filtered to retain reads containing the expected transposon-end sequence. Filtered reads were aligned to the corresponding target sequence using Geneious. The integration position was calculated as the distance between the 3′ edge of the crRNA target site and the 5′ end of the transposon insertion.

### Quantitative PCR analysis of integration efficiency

CAST integration efficiency was quantified using a duplex probe-based quantitative PCR (qPCR) assay in which the integration junction and genomic reference locus were amplified in the same reaction. For each 10 µL reaction, 5 µL of 2× Luna® Universal Probe qPCR Master Mix (NEB, catalog no. M3004S), 0.4 µL each of 10 µM forward and reverse primers targeting the integration junction, 0.2 µL of 10 µM FAM-labeled junction probe, 0.4 µL each of 10 µM forward and reverse primers targeting the *NbRDR1* reference gene, 0.2 µL of 10 µM HEX-labeled reference probe, 2 µL of genomic DNA template, and 1 µL of nuclease-free water were combined. Oligonucleotide sequences were synthesized by Integrated DNA Technologies and are listed in **Supplementary Table 4**. Reactions were assembled in 96-well qPCR plates, sealed, briefly centrifuged, and run on Bio-Rad CFX96 real-time PCR detection system using probe-detection settings appropriate for FAM- and HEX-labeled probes. Thermocycling was performed under the following conditions: initial denaturation at 95 °C for 60 s; followed by 40 cycles of denaturation at 95 °C for 15 s and extension at 60 °C for 30 s, with plate read during the 60 °C extension step. Integration efficiency was calculated by normalizing the FAM-labeled integration-junction signal to the HEX-labeled reference-gene signal using the ΔCq method, as 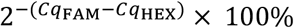.

### Droplet digital PCR analysis of integration efficiency

Duplex 22 µl ddPCR reactions were prepared by mixing 11 µl of ddPCR Supermix for Probes (no dUTP; Bio-Rad, catalog no. 1863024), 900 nM forward and reverse primers for the target and reference loci, 250 nM HEX-labeled probe targeting the *NbRDR1* reference gene, 250 nM FAM-labeled probe targeting the integrated junction, and 50-100 ng of genomic DNA. Oligonucleotide sequences were synthesized by Integrated DNA Technologies and are listed in **Supplementary Table 4**. Reaction mixtures were loaded into a DG8 cartridge (Bio-Rad, catalog no. 1864007) along with 70 µl of droplet generation oil (Bio-Rad, catalog no. 1863005), and droplets were generated using a Bio-Rad QX200 Droplet Generator. Following droplet generation, 40 µl of emulsion was transferred to a 96-well plate, heat-sealed with pierceable foil, and thermal-cycled under the manufacturer’s conditions with an annealing/extension temperature of 58°C. Droplets were read using a Bio-Rad QX200 Droplet Reader. Data were analyzed using QX Manager software (Bio-Rad). Integration efficiency was calculated as the ratio of FAM-positive integration-junction copies to HEX-positive reference-locus copies.

### Candidate cofactor screen

Candidate cofactors were screened to identify proteins that modulate evoCAST integration efficiency in plant cells. The screened panel included EcClpP, ScDDR2, ScRTT10, AtHMGB2, AtHMGB3, EcRecA, ScRad54, and AtRad54. Coding sequences were codon-optimized for expression in *N. benthamiana* using the GenSmart™ Codon Optimization tool, expressed under the control of the pUBQ10 promoter and tHSP terminator, and are listed in **Supplementary Table 5**. Each candidate cofactor was co-delivered with the rest of evoCAST components by *Agrobacterium*-mediated infiltration in *N. benthamiana*. Integration efficiency was quantified by qPCR and expressed as fold change relative to the evoCAST + ClpX baseline condition lacking an additional cofactor.

## Data Availability Statement

All data are included either in the manuscript or in the supplementary files.

## Author Contributions

Conceptualization: YW, GSD

Methodology: YW, KTM

Data Acquisition: YW, KTM, MGL, EL, AP

Supervision: GSD

Writing: YW, KTM, GSD

Funding Acquisition: GSD

## Acknowledgements

The authors thank Professor Carlotta Ronda (Innovative Genomics Institute), Isaac P. Witte (Harvard University), George D. Lampe (Columbia University), Professor Kaihang Wang (Caltech), and Jolena Zhou (Caltech) for valuable discussions and feedback. The authors acknowledge the Caltech Biological Imaging Facility, with support from the Caltech Beckman Institute and the Arnold and Mabel Beckman Foundation, for access to the Leica STELLARIS 8 inverted confocal microscope. The authors also acknowledge the Beckman Institute CLOVER Center for access to droplet digital PCR instrumentation and experimental support. Schematics were created with BioRender.com.

## References

1. Gao, C. Genome engineering for crop improvement and future agriculture. Cell 184, 1621–1635 (2021).

2. Zhu, H., Li, C. & Gao, C. Applications of CRISPR–Cas in agriculture and plant biotechnology. Nat Rev Mol Cell Biol 21, 661–677 (2020).

3. Wang, Y. et al. Simultaneous editing of three homoeoalleles in hexaploid bread wheat confers heritable resistance to powdery mildew. Nat Biotechnol 32, 947–951 (2014).

4. He, F. et al. Simultaneous editing of three homoeologues of TaCIPK14 confers broad-spectrum resistance to stripe rust in wheat. Plant Biotechnology Journal 21, 354–368 (2023).

5. Su, H. et al. Generation of the transgene-free canker-resistant Citrus sinensis using Cas12a/crRNA ribonucleoprotein in the T0 generation. Nat Commun 14, 3957 (2023).

6. Fan, T. et al. High performance TadA-8e derived cytosine and dual base editors with undetectable o[-target e[ects in plants. Nat Commun 15, 5103 (2024).

7. Molla, K. A., Sretenovic, S., Bansal, K. C. & Qi, Y. Precise plant genome editing using base editors and prime editors. Nat. Plants 7, 1166–1187 (2021).

8. Zong, Y. et al. An engineered prime editor with enhanced editing efficiency in plants. Nat Biotechnol 40, 1394–1402 (2022).

9. Jiang, Y. et al. Optimized prime editing efficiently generates glyphosate-resistant rice plants carrying homozygous TAP-IVS mutation in EPSPS. Molecular Plant 15, 1646–1649 (2022).

10. Vu, T. V. et al. Optimized dicot prime editing enables heritable desired edits in tomato and Arabidopsis. Nat. Plants 10, 1502–1513 (2024).

11. Karmakar, S. et al. A miniature alternative to Cas9 and Cas12: Transposon-associated TnpB mediates targeted genome editing in plants. Plant Biotechnology Journal 22, 2950–2953 (2024).

12. Zhu, J. et al. Engineering hypercompact IscB nucleases for efficient and versatile genome editing in rice. Genome Biol 27, 49 (2026).

13. Li, Q. et al. Genome editing in plants using the TnpB transposase system. aBIOTECH 5, 225–230 (2024).

14. Weiss, T. et al. Viral delivery of an RNA-guided genome editor for transgene-free germline editing in Arabidopsis. Nat. Plants 11, 967–976 (2025).

15. Dong, O. X. & Ronald, P. C. Targeted DNA insertion in plants. Proceedings of the National Academy of Sciences 118, e2004834117 (2021).

16. Vollen, K., Alonso, J. M. & Stepanova, A. N. Beyond a few bases: methods for large DNA insertion and gene targeting in plants. The Plant Journal 121, e70099 (2025).

17. Agrobacterium-Mediated Plant Transformation: the Biology behind the “Gene-Jockeying” Tool | Microbiology and Molecular Biology Reviews. 10.1128/mmbr.67.1.16-37.2003?url_ver=Z39.88-2003&rfr_id=ori%3Arid%3Acrossref.org&rfr_dat=cr_pub++0pubmed.

18. Dahan-Meir, T. et al. Efficient in planta gene targeting in tomato using geminiviral replicons and the CRISPR/Cas9 system. The Plant Journal 95, 5–16 (2018).

19. Zhou, Z. et al. Cas9-Rep fusion tethers donor DNA in vivo and boosts the efficiency of HDR-mediated genome editing. Plant Biotechnology Journal 23, 2006–2017 (2025).

20. Ellison, E. E. et al. Multiplexed heritable gene editing using RNA viruses and mobile single guide RNAs. Nat. Plants 6, 620–624 (2020).

21. Lu, Y. et al. Targeted, efficient sequence insertion and replacement in rice. Nat Biotechnol 38, 1402–1407 (2020).

22. Li, S. et al. Precise gene replacement in rice by RNA transcript-templated homologous recombination. Nat Biotechnol 37, 445–450 (2019).

23. Sun, C. et al. Precise integration of large DNA sequences in plant genomes using PrimeRoot editors. Nat Biotechnol 42, 316–327 (2024).

24. Liu, P. et al. Transposase-assisted target-site integration for efficient plant genome engineering. Nature 631, 593–600 (2024).

25. Wei, S. et al. Transposase-Assisted Donor Tethering Boosts Large-Fragment HDR in Plants. Advanced Science 11, e75565.

26. Muchenje, K. T. et al. Optimized R2 retroelement complexes for DNA insertion into plant genomes. Nat Biotechnol 1–13 (2026) doi:10.1038/s41587-026-03197-y.

27. Strecker, J. et al. RNA-guided DNA insertion with CRISPR-associated transposases. Science 365, 48–53 (2019).

28. Klompe, S. E., Vo, P. L. H., Halpin-Healy, T. S. & Sternberg, S. H. Transposon-encoded CRISPR–Cas systems direct RNA-guided DNA integration. Nature 571, 219–225 (2019).

29. Halpin-Healy, T. S., Klompe, S. E., Sternberg, S. H. & Fernández, I. S. Structural basis of DNA targeting by a transposon-encoded CRISPR–Cas system. Nature 577, 271–274 (2020).

30. Finocchio, G., Querques, I., Chanez, C., Speichert, K. J. & Jinek, M. Structural basis of TnsC oligomerization and transposase recruitment in type I-B CRISPR-associated transposons. Nucleic Acids Research 53, gkaf149 (2025).

31. Walker, M. W. G., Klompe, S. E., Zhang, D. J. & Sternberg, S. H. Novel molecular requirements for CRISPR RNA-guided transposition. Nucleic Acids Research 51, 4519–4535 (2023).

32. Petassi, M. T., Hsieh, S.-C. & Peters, J. E. Guide RNA Categorization Enables Target Site Choice in Tn7-CRISPR-Cas Transposons. Cell 183, 1757–1771.e18 (2020).

33. Klompe, S. E. et al. Evolutionary and mechanistic diversity of Type I-F CRISPR-associated transposons. Molecular Cell 82, 616–628.e5 (2022).

34. Vo, P. L. H. et al. CRISPR RNA-guided integrases for high-efficiency, multiplexed bacterial genome engineering. Nat Biotechnol 39, 480–489 (2021).

35. Lampe, G. D. et al. Targeted DNA integration in human cells without double-strand breaks using CRISPR-associated transposases. Nat Biotechnol https://doi.org/10.1038/s41587-023-01748-1 (2023) doi:10.1038/s41587-023-01748-1.

36. Witte, I. P. et al. Programmable gene insertion in human cells with a laboratory-evolved CRISPR-associated transposase. Science 388, eadt5199 (2025).

37. Philips J. G. et al. The widely used Nicotiana benthamiana 16c line has an unusual T-DNA integration pattern including a transposon sequence. PLOS ONE 12, e0171311 (2017).

38. Liu, Z. et al. Systematic comparison of 2A peptides for cloning multi-genes in a polycistronic vector. Sci Rep 7, 2193 (2017).

39. Ke, Y. et al. Precise knock-in of stress-responsive cis-regulatory elements using gene targeting for improving abiotic stress tolerance in plants. New Phytologist 247, 2147–2162 (2025).

40. Baker, T. A. & Sauer, R. T. ClpXP, an ATP-powered unfolding and protein-degradation machine. Biochimica et Biophysica Acta (BBA) - Molecular Cell Research 1823, 15–28 (2012).

41. Zhou, X. et al. Artificial optimization of bamboo Ppmar2 transposase and host factors effects on Ppmar2 transposition in yeast. Front. Plant Sci. 13, (2022).

42. Zayed, H., Izsvák, Z., Khare, D., Heinemann, U. & Ivics, Z. The DNA-bending protein HMGB1 is a cellular cofactor of Sleeping Beauty transposition. Nucleic Acids Research 31, 2313–2322 (2003).

43. Grasser, K. D. Chromatin-associated HMGA and HMGB proteins: versatile co-regulators of DNA-dependent processes. Plant Mol Biol 53, 281–295 (2003).

44. Song, L. C. T. et al. Identification of proteins influencing CRISPR-associated transposases for enhanced genome editing. Science Advances 12, eaea1429 (2026).

45. Shaked, H., Melamed-Bessudo, C. & Levy, A. A. High-frequency gene targeting in Arabidopsis plants expressing the yeast RAD54 gene. Proceedings of the National Academy of Sciences 102, 12265–12269 (2005).

46. Osakabe, K. et al. Isolation and characterization of the RAD54 gene from Arabidopsis thaliana. The Plant Journal 48, 827–842 (2006).

47. George, J. T. et al. Mechanism of target site selection by type V-K CRISPR-associated transposases. Science 382, eadj8543 (2023).

48. Tou, C. J., Orr, B. & Kleinstiver, B. P. Precise cut-and-paste DNA insertion using engineered type V-K CRISPR-associated transposases. Nat Biotechnol 41, 968–979 (2023).

49. Stoddard, B. L. Homing endonucleases from mobile group I introns: discovery to genome engineering. Mobile DNA 5, 7 (2014).

50. Engineered geminivirus replicons enable rapid in planta directed evolution | Science. https://www.science.org/doi/10.1126/science.ady2167.

51. Levchenko, I., Luo, L. & Baker, T. A. Disassembly of the Mu transposase tetramer by the ClpX chaperone. Genes Dev. 9, 2399–2408 (1995).

52. Holder, J. W. & Craig, N. L. Architecture of the Tn7 Posttransposition Complex: An Elaborate Nucleoprotein Structure. Journal of Molecular Biology 401, 167–181 (2010).

53. Witte, I. P. et al. Programmable gene insertion in human cells with a laboratory-evolved CRISPR-associated transposase. Science 388, eadt5199 (2025).

54. Weiss, T. et al. Epigenetic features drastically impact CRISPR–Cas9 efficacy in plants. Plant Physiol 190, 1153–1164 (2022).

55. Carlson, E. D., Rajniak, J. & Sattely, E. S. Multiplicity of the Agrobacterium Infection of Nicotiana benthamiana for Transient DNA Delivery. ACS Synth. Biol. 12, 2329–2338 (2023).

56. Gelvin, S. B., Singer, K., Lee, L.-Y. & Yuan, J. Characterization of T-Circles and Their Formation Reveal Similarities to Agrobacterium T-DNA Integration Patterns. Front. Plant Sci. 13, (2022).

57. Pedersen, D. S. & Grasser, K. D. The role of chromosomal HMGB proteins in plants. Biochimica et Biophysica Acta (BBA) - Gene Regulatory Mechanisms 1799, 171–174 (2010).

58. Yoo, S.-D., Cho, Y.-H. & Sheen, J. Arabidopsis mesophyll protoplasts: a versatile cell system for transient gene expression analysis. Nat Protoc 2, 1565–1572 (2007).

