## Supplementary Figures for "CRISPR-Associated Transposases Enable Programmable DNA Integration in Plants"

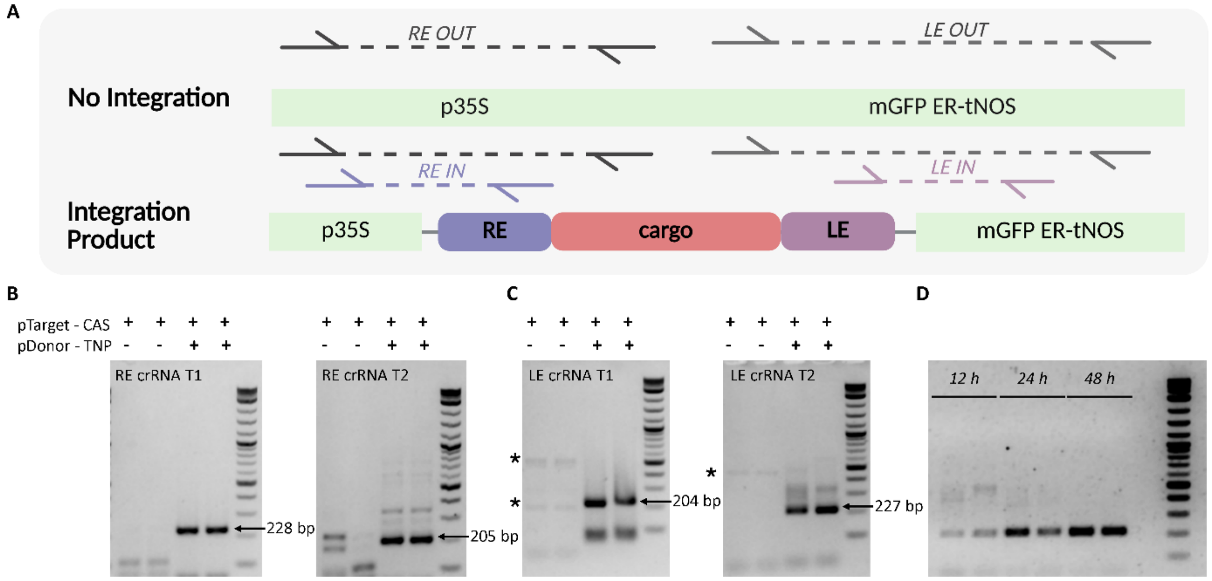


**Supplementary Figure 1. Nested PCR detection of PseCAST-mediated episomal integration in *A. thaliana* protoplasts. (A)** Schematic of the nested PCR design. Upon CAST-mediated insertion of the RE-cargo-LE transposon between the p35S promoter and mGFP ER-tNOS reporter cassette, junction-specific primer pairs amplify integration products at the right-end (RE) and left-end (LE) junctions. **(B)** Nested junction PCR detecting RE integration products for crRNA T1 and crRNA T2 in *A. thaliana* protoplasts co-transfected with pTarget-CAS and pDonor-TNP. Bands of the expected sizes are indicated by arrows. **(C)** Nested junction PCR detecting LE integration products for crRNA T1 and crRNA T2 in *A. thaliana* protoplasts co-transfected with pTarget-CAS and pDonor-TNP. Bands of the expected sizes are indicated by arrows. Asterisks indicate nonspecific PCR amplification bands. **(D)** Time-course analysis of episomal integration products detected at 12, 24, and 48 h after protoplast transfection. p35s: CaMV 35S promoter, mGFP ER: endoplasmic reticulum-targeted modified green fluorescent protein, tNOS: nopaline synthase terminator.

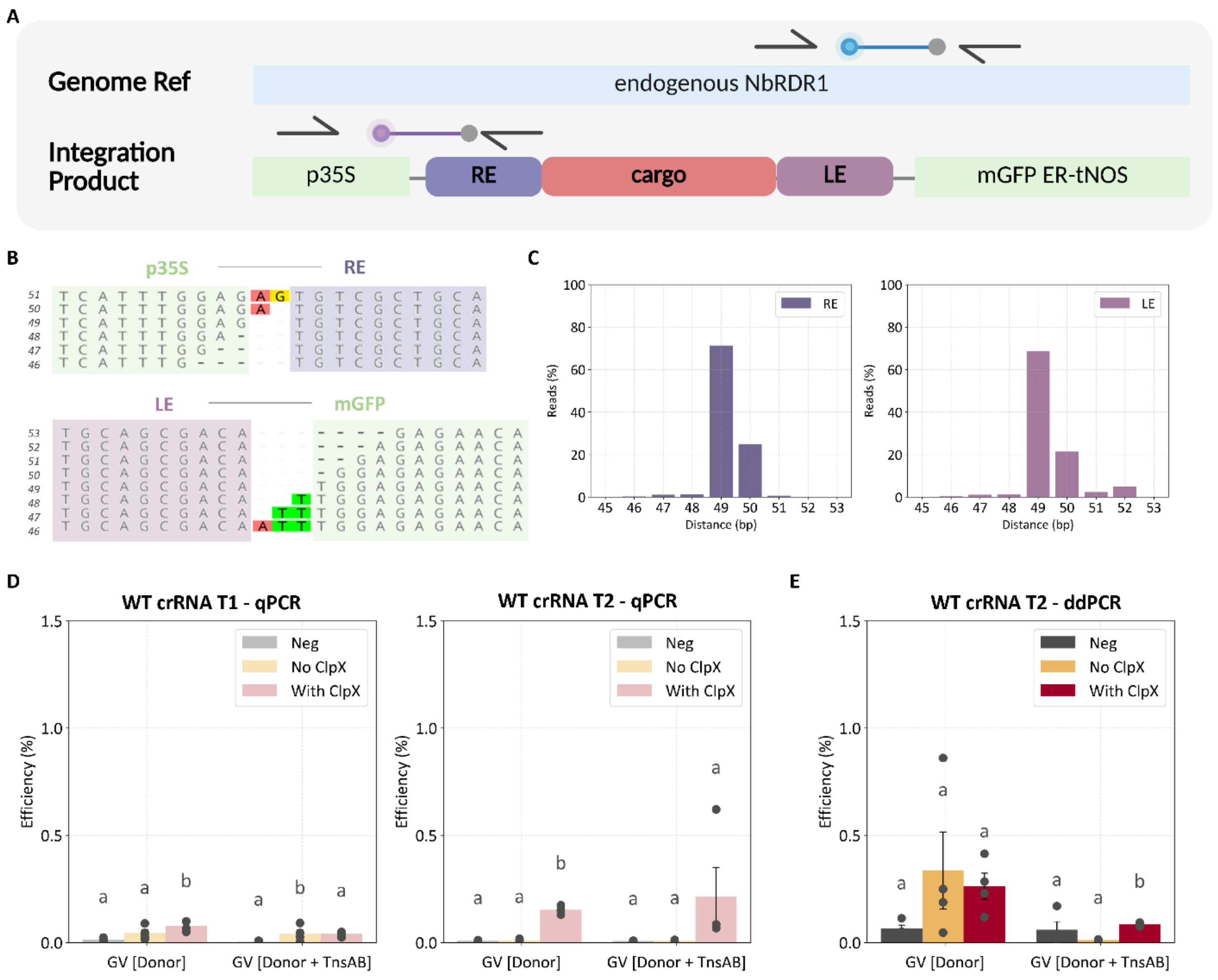


**Supplementary Figure 2. Quantification and sequence analysis of wild-type PseCAST integration in 16C *N. benthamiana*. (A)** Schematic of the probe-based quantitative PCR (qPCR) and droplet digital PCR (ddPCR) assays used to quantify chromosomal integration. Primers and probe targeting the RE junction of the integrated product were used to quantify integration frequency, normalized to the endogenous NbRDR1 reference locus. **(B)** Representative NGS reads spanning the p35S-RE junction (top) and LE-mGFP junction (bottom). **(C)** Distribution of insertion distances at the RE (left) and LE (right) junctions. **(D)** Integration efficiency measured by qPCR for wild-type PseCAST using crRNA T1 (left) and crRNA T2 (right) across GV-Donor and GV-Donor+TnsAB configurations, with or without ClpX co-expression. **(E)** Integration efficiency measured by ddPCR for wild-type PseCAST using crRNA T2 across GV-Donor and GV-Donor+TnsAB configurations, with or without ClpX co-expression. Bars represent mean ± S.E.M.; individual points indicate biological replicates (n=4). Statistical analysis was performed using one-way ANOVA with Tukey's HSD post hoc test. Different letters indicate statistically significant differences ($p < 0.05$). NbRDR1: *N. benthamiana* RNA-dependent RNA polymerase 1, GV: geminiviral replicon, p35s: CaMV 35S promoter, mGFP ER: endoplasmic reticulum-targeted modified green fluorescent protein, tNOS: nopaline synthase terminator, WT: wild-type.

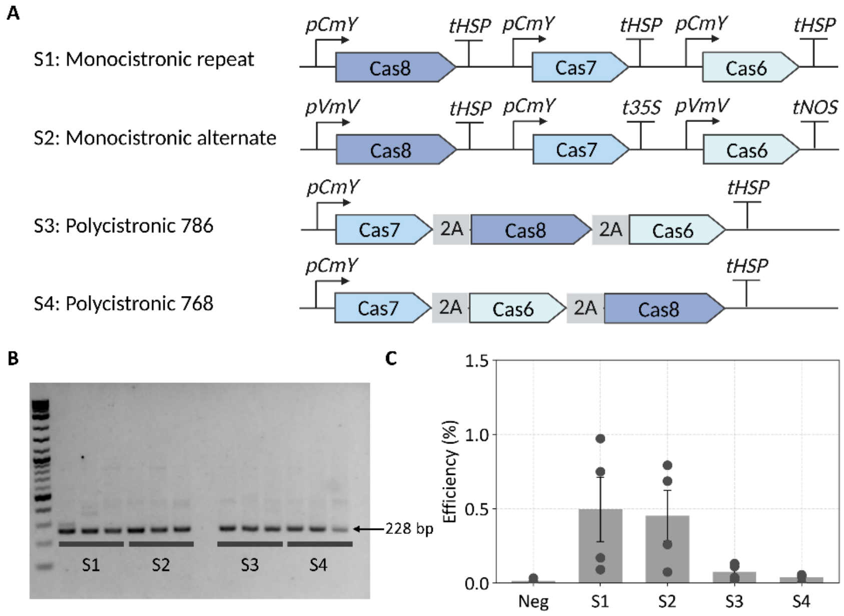


**Supplementary Figure 3. Comparison of Cascade protein expression architectures for CAST activity in *N. benthamiana*. (A)** Schematic of four Cascade expression architectures tested for wild-type PseCAST activity. S1 contains monocistronic Cas8, Cas7, and Cas6 expression cassettes driven by repeated pCmY promoters and tHSP terminators. S2 contains monocistronic cassettes with alternating promoter and terminator elements. S3 and S4 contain 2A-linked polycistronic Cascade designs with different Cas7, Cas8, and Cas6 gene orders. **(B)** Nested junction PCR detecting CAST-mediated integration products for the indicated Cascade expression architectures. The expected 228-bp product is indicated by an arrow. **(C)** Integration efficiency of the indicated Cascade expression architectures measured by qPCR. Bars represent mean ± S.E.M.; individual points indicate biological replicates (n=4). No statistically significant differences were detected among the tested Cascade architectures after one-way ANOVA with Tukey's HSD post hoc test ($p > 0.05$). pCmY: CmYLCV6 promoter, tHSP: HSP18.2 terminator, pVmv: Cassava vein mosaic virus promoter, p35s: CaMV 35S promoter, t35s: CaMV 35S terminator, tNOS: nopaline synthase terminator, 2A: porcine teschovirus-1 2A peptide.

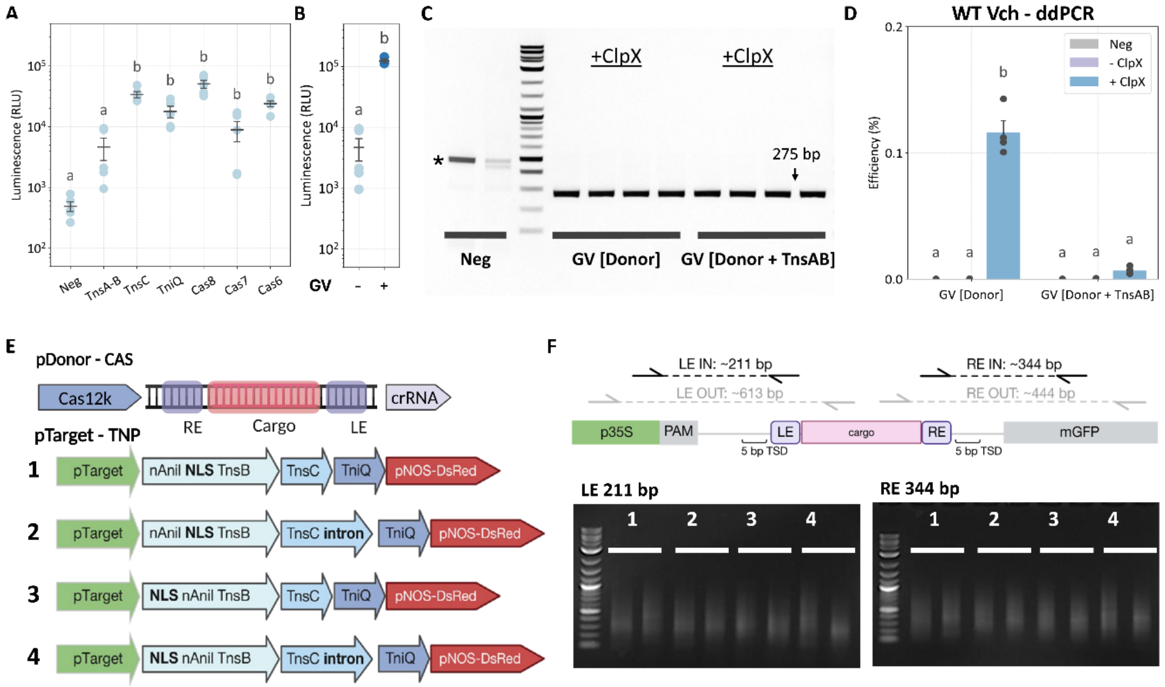


**Supplementary Figure 4. Characterization of alternative CAST systems. (A)** Luminescence-based detection of individual HiBiT-tagged VchCAST components in plant nuclear fractions. **(B)** Geminiviral replicon (GV)-mediated enhancement of TnsA-TnsB detection in plant nuclear fractions. Bars represent mean ± S.E.M.; individual points indicate biological replicates (n=5). **(C)** VchCAST chromosomal integration assay in 16C *N. benthamiana*. VchCAST components were delivered using the GV-Donor or GV-Donor+TnsAB configurations. Nested PCR detected the integrated junction in ClpX-containing conditions. Bands of the expected sizes are indicated by arrows. Asterisks indicate nonspecific PCR amplification bands. **(D)** Integration efficiency measured by ddPCR for VchCAST across GV-Donor and GV-Donor+TnsAB configurations, with or without ClpX co-expression. Bars represent mean ± S.E.M.; individual points indicate biological replicates (n=4). Statistical analysis was performed using one-way ANOVA with Tukey's HSD post hoc test. Different letters indicate statistically significant differences ($p < 0.05$). **(E)** Schematic of the Type V-K CAST construct design. The pDonor-CAS construct encodes the Cas12k, the crRNA, and the mini-transposon cargo. The pTarget-TNP constructs encode the target site and HELIX-derived transposition machinery. Four transposition machinery configurations were tested, differing in NLS and I-AniI placement and intron usage. **(F)** Junction PCR assay design and representative gel analysis for detecting Type V-K CAST integration at the LE and RE junctions. Expected LE and RE integration products are approximately 211 bp and 344 bp, respectively. No clear junction products of the expected sizes were detected. VchCAST: *Vibrio cholerae* CRISPR-associated transposase, p35s: CaMV 35S promoter, mGFP ER: endoplasmic reticulum-targeted modified green fluorescent protein, tNOS: nopaline synthase terminator, NLS: nuclear localization signal.

A

| Component | Optimization method | Change |
|---|---|---|
| **TnsB** | PACE | F43S, Y349N, P352T, A390V, D396N, Q410K, H464R, V526E, Q549R, Q594L |
| **TnsA** | PACE | P88T, I147V, V170L, F180L, F182L |
| **TnsC** | PACE | R197I, A314K |
| **Cas7** | Rational engineering | N347K (DNA contact) |
| **Cas8** | Rational engineering | N125D, A244N, A410R (PAM recognition) + N terminal 2x bipartite NLS |
| **Cas6** | Rational engineering | N terminal 2x bipartite NLS |
| **TniQ** | Rational engineering | N terminal 2x bipartite NLS |

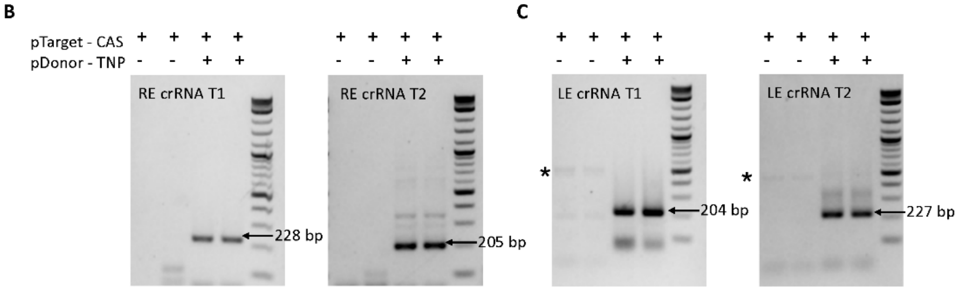


**Supplementary Figure 5. EvoCAST component composition and episomal integration activity in *A. thaliana* protoplasts. (A)** Summary of evoCAST modifications relative to wild-type PseCAST. PACE-derived substitutions are present in TnsB, TnsA, and TnsC, while rationally engineered changes include a Cas7 DNA-contact substitution, Cas8 PAM-recognition substitutions, and additional N-terminal bipartite nuclear localization signals on Cas8, Cas6, and TniQ[1]. **(B)** Nested junction PCR detecting RE integration products for crRNA T1 and crRNA T2 in *A. thaliana* protoplasts co-transfected with pTarget-CAS and pDonor-TNP. Bands of the expected sizes are indicated by arrows. **(C)** Nested junction PCR detecting LE integration products for crRNA T1 and crRNA T2 in *A. thaliana* protoplasts co-transfected with pTarget-CAS and pDonor-TNP. Bands of the expected sizes are indicated by arrows. Asterisks indicate nonspecific PCR amplification bands. PACE: phage-assisted continuous evolution, NLS: nuclear localization signal, PAM: protospacer adjacent motif.

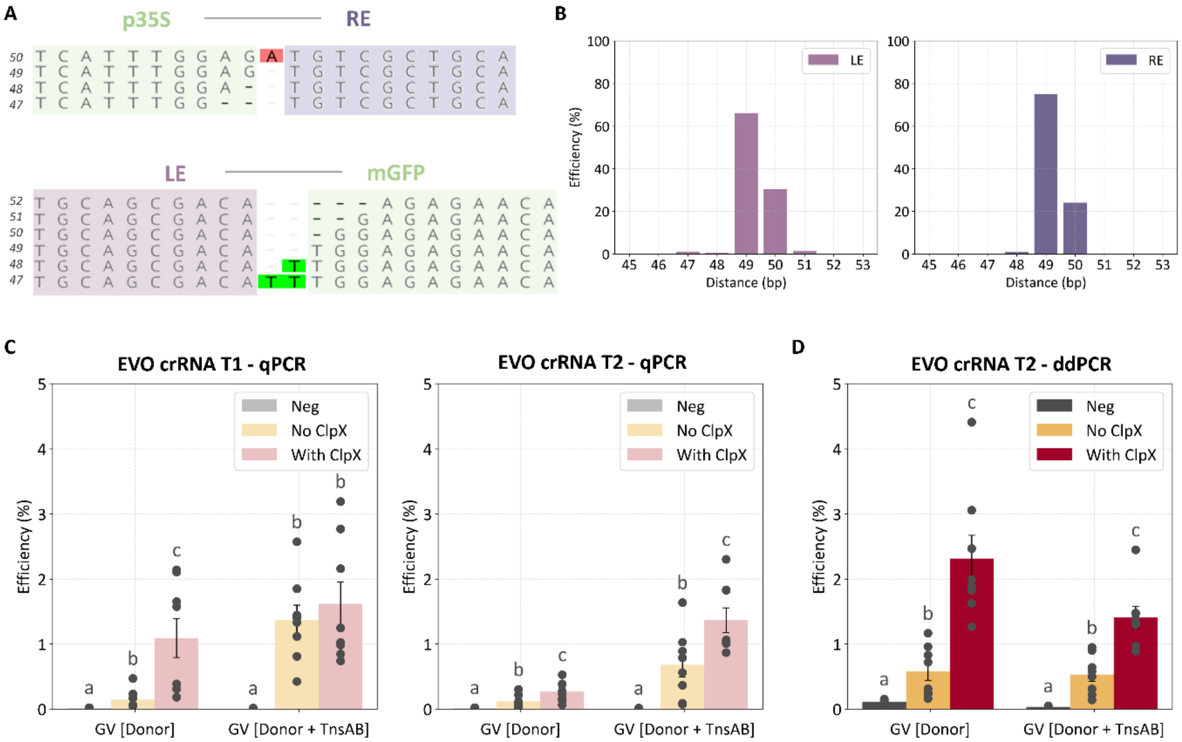


**Supplementary Figure 6. Quantification and sequence analysis of evoCAST integration in 16C *N. benthamiana*. (A)** Representative NGS reads spanning the p35S-RE junction (top) and LE-mGFP junction (bottom). **(B)** Distribution of insertion distances at the RE (left) and LE (right) junctions. **(C)** Integration efficiency measured by qPCR for evoCAST using crRNA T1 (left) and crRNA T2 (right) across GV-Donor and GV-Donor+TnsAB configurations, with or without ClpX co-expression. **(D)** Integration efficiency measured by ddPCR for evoCAST using crRNA T2 across GV-Donor and GV-Donor+TnsAB configurations, with or without ClpX co-expression. Bars represent mean ± S.E.M.; individual points indicate biological replicates (n=8). Statistical analysis was performed using one-way ANOVA with Tukey's HSD post hoc test. Different letters indicate statistically significant differences ($p < 0.05$). GV: geminiviral replicon.

### References


1. Witte, I. P. *et al.* Programmable gene insertion in human cells with a laboratory-evolved CRISPR-associated transposase. *Science* **388**, eadt5199 (2025).
